# Detection of AmpC *β*-lactamases in *Escherichia coli* using different screening agars

**DOI:** 10.1101/787085

**Authors:** Evert den Drijver, Jaco J. Verweij, Carlo Verhulst, Joke Soer, Kees Veldman, John W. Rossen, A.M.D. (Mirjam) Kooistra-Smid, Marjolein F. Q. Kluytmans van den Bergh, Jan A. J. W. Kluytmans

## Abstract

The aim of this study was to determine the performance of both cefotaxime and ceftazidime containing agars on the specificity and sensitivity for chromosomal AmpC-hyperproducing and plasmid AmpC harboring *Escherichia coli* compared to ESBL-producing *E. coli* and *E. coli* without ESBL, pAmpC or cAmpC hyperproduction. Second, we evaluated the influence of adding cefoxitin to these agars for detection of both chromosomal AmpC-hyperproducing and plasmid AmpC harboring *E. coli.*

Four different homemade screening agars with cefotaxime (1mg/L), ceftazidime (1mg/L), cefotaxime (1mg/L) with cefoxitin (8mg/L), and ceftazidime (1mg/L) with cefoxitin (8mg/L) were compared to each other for the identification of AmpC producing *E. coli.* A total of 40 isolates with plasmid encoded AmpC *β*-lactamases, 40 isolates with alterations in the promoter/attenuator region of the AmpC gene leading to hyperproduction of the *β*-lactamase, 40 isolates with ESBL genes and 39 isolates lacking both a AmpC and ESBL genotype were used to test the four agars.

The sensitivity and specificity were 100% (95% confidence interval (95% CI) 96.1% to 100%) and 48.1% (95% CI 38.6%-60.2%), respectively, for the cefotaxime agar; 100% (95% CI 96.1% to 100%) and 49.41% (95% CI 39.8%-61.4%), respectively, for the ceftazidime agar; 96.3% (95% CI 89.1% to 99.2%) and 77.2% (95% CI 66.7%-85.2%) respectively, for the cefotaxime with cefoxitin agar; 98.8% (95% CI) 92.6% to 99.6%) and 81.0% (95% CI 70.9%-88.3%) respectively, for the ceftazidime agar with cefoxitin. The main reason for false-positive results were ESBL-harboring strains that grew on various agars; therefore, the specificity of each agar reported here was influenced mainly by the proportion of ESBL isolates tested. In conclusion addition of cefoxitin to cefotaxime and ceftazidime containing agars had little influence on sensitivity, but increased specificity for the detection of AmpC in *E. coli*.

## Introduction

Over the past decades, the importance of resistant Enterobacteriales increased substantially due to the emergence of various resistance traits that inactivate most β-lactam antibiotics. In *Escherichia coli*, resistance to 2^nd^ and 3^rd^ generation cephalosporins is most frequently due to the occurrence of plasmid-encoded ESBL, as well as both plasmid-encoded (*pampC*) and chromosomally-encoded (*campC*) AmpC genes. As the rapid and accurate detection of resistant bacteria is crucial for treatment and control, selective media for detection of extended-spectrum *β*-lactamase-producing Enterobacteriales (ESBL-E) have been developed. A considerable number of studies were published on the performance of ESBL-E (1–5) screening agars. So far, few studies have focused on screening media for AmpC *β*-lactamase-producing Enterobacteriales (AmpC-E).

The basis of most agars that are selective for ESBL is a adding a 3^rd^ generation cephalosporin, e.g. cefotaxime or ceftazidime. Some ESBL-E screening agars use AmpC inhibitors, such as cloxacillin (3–5) to suppress Enterobacteriales that intrinsically produce AmpC *β*-lactamase on high level (e.g. *Enterobacter* spp). Overgrowth of such species with intrinsic AmpC *β*-lactamase hyperproduction makes it more difficult to screen for *E. coli* with acquired *β*-lactamases, which may lead to a lower sensitivity of the ESBL-E screening agars(3). *E. coli* isolates produce chromosomally encoded AmpC (cAmpC) constitutively, but only on a low level. However, they may acquire alterations in the promoter/attenuator region leading to hyperproduction of cAmpC *β*-lactamase (6,7). Growth of both cAmpC and pAmpC producing Enterobacteriales is inhibited by cloxacillin. Cloxacillin containing agars may improve the yield of ESBL producing *E. coli* but they lead to a lower yield of both pAmpC and cAmpC *β*-lactamase producing *E. coli*. One option to solve this problem could be to include an additional agar specifically developed for AmpC producing *E. coli* during screening.

According to the EUCAST guidelines combined resistance to 3^rd^ generation cephalosporins and cefoxitin, a cephamycin antibiotic, may be used as phenotypic criteria for AmpC screening (8). Cefoxitin is highly active against ESBL producing bacteria (9), but has little activity against most AmpC producing bacteria (10). This makes cefoxitin useful in screening strategies for AmpC (11–13). A study by Reuland *et al* compared three different screening strategies of pAmpC Enterobacteriales (14) based on susceptibility patterns. These included: reduced susceptibility to cefotaxime and/or ceftazidime (>1 mg/L), reduced susceptibility to cefoxitin (> 8mg/L), or a combination of cefotaxime and/or ceftazidime (>1 mg/L). Using just cefoxitin during a screening resulted in high sensitivity (97%), but was not specific for AmpC (72%) as a porin deficiency can cause cefoxitin resistance as well. Combining reduced cefotaxime and/or ceftazidime susceptibility with reduced cefoxitin susceptibility increased the specificity to 90%, without significant loss of sensitivity (97%). Reuland et al. did not use a screening agar to differentiate pAmpC Enterobacteriales. We hypothesize that an agar which combines cefoxitin with either cefotaxime or ceftazidime may result in an AmpC specific screening agar, which could be of additional value to the use of an ESBL-agar with cloxacillin.

The first aim of this study was to determine the performance of both cefotaxime and ceftazidime containing agars on the specificity and sensitivity for AmpC-producing *E. coli* compared to ESBL-producing and AmpC/ESBL negative *E. coli*. Second, we evaluated the influence of adding cefoxitin to these agars for detection of AmpC producing *E. coli*.

## Methods

### Collection of isolates

A panel of 159 *E. coli* isolates was used for the evaluation of the AmpC screening agar. This panel consisted of 40 *E. coli* isolates harboring pAmpC, 40 *E. coli* isolates with cAmpC mutations known to lead to hyperproduction (referred in this study as “cAmpC positive”), 40 ESBL-producing *E. coli* isolates and 39 *E. coli* without signs of ESBL, pAmpC or cAmpC hyperproduction based on genotype (referred in this study as “AmpC/ESBL negative”). The ESBL-producing *E. coli* and AmpC/ESBL negative *E. coli* isolates were confirmed to lack any known alterations in the promoter/attenuator region of the AmpC gene leading to hyperproduction of the *β*-lactamase, as reported by Tracz *et al* (7). Isolates were recovered at different study sites or from various clinical specimens, and were isolated from humans (n=141) and animals (n=18) (15–20). More detailed information on the recovery of the isolates is described in the additional file 1. Additional file 2 shows an overview of all strains.

### Evaluation of AmpC agars

MacConkey agar no3 medium was obtained from Oxoid Limited (ThermoFisher, Basingstoke, Hampshire, United Kingdom). Both cefotaxime sodium salt and ceftazidime sodium salt were obtained from VWR International (Radnor, Pennsylvania, USA). Cefoxitin sodium salt was obtained from Sigma-Aldrich (Poole, Dorset, UK). MacConkey agar no3 was used as the basal medium to which different dilutions of antibiotics were added. Four different concentration combination were evaluated; 1. cefotaxime 1mg/L agar, 2. ceftazidime 1 mg/L agar, 3. cefotaxime 1mg/L with cefoxitin 8mg/L agar, and 4. ceftazidime 1 mg/L with cefoxitin 8mg/L agar.

All strains with AmpC or ESBL were cultured in a selective enrichment broth consisting of a TSB containing 0, 25 mg/L cefotaxime and 8mg/L vancomycin (TSB-VC, Mediaproducts, Groningen, The Netherlands). For strains without AmpC or ESBL a brain heart infusion broth was used (BHI, Mediaproducts, Groningen, The Netherlands), as using selective enrichment broth might inhibit growth. After overnight aerobic incubation for 18-24 hours (35°-37°), 10 µl of the broth was subcultured on the four different AmpC agars. Agars were incubated aerobically for 18-24 hours (35°-37°). Results were interpreted as growth or no growth.

### Statistical analysis

Data were analyzed with Statistical Package for Social Science software (SPSS; IBM Corp., Armonk, New York, US; version 22). For sensitivity all strains positive for an acquired AmpC gene or alterations in promoter/attenuator AmpC related to hyperproduction were considered true positives. For specificity were all strains negative for an acquired AmpC gene and alterations in promoter/attenuator AmpC related to hyperproduction were considered true negatives. Confidence intervals (95% CI) were calculated according to the adjusted Wald method.

The prevalence of ESBL in the panel of isolates was 25.16% (40 out of 159 isolates), due to the artificial nature of the collection set up. To analyse the influence of variations in ESBL prevalence on the specificity of both the cefotaxime 1mg/L + cefoxitin 8mg/L and the ceftazidime 1mg/L + cefoxitin 8mg/L agar, we remodelled our data for an ESBL prevalence range from 0 to 99%.

Specificity of the cefotaxime 1mg/L + cefoxitin 8mg/L agar was calculated using following formula: 38 + (23/40 x number of ESBL isolates) / (38 + (23/40 x number of ESBL isolates) + (1 + 17/40 x number of ESBL isolates)).

Specificity of the ceftazidime 1mg/L + cefoxitin 8mg/L agar was calculated using following formula: 39 + (25/40 x number of ESBL isolates)/(39 + (25/40 x number of ESBL isolates) + (15/40 x number of ESBL isolates)). Prevalence of ESBL was calculated using the formula: number of ESBL isolates / (119 + number of ESBL isolates).

## Results

The 159 *E. coli* isolates were evaluated for growth on the four different selective agar plates as shown in Table 1. Isolates hyperproducing cAmpC or producing pAmpC or ESBL *E. coli* grew on both cefotaxime and ceftazidime agar plates without cefoxitin. Of the AmpC/ESBL negative isolates, only one (2.6%) isolate grew on the cefotaxime agar. No growth of AmpC/ESBL negative isolates was seen on the ceftazidime agar.

**TABLE 1.** Ability of 159 E. coli isolates to grow on four different AmpC screening agars

| Characteristics | No. of isolates | Cefotaxime 1 mg/L |  | Ceftazidime 1 mg/L |  |
| --- | --- | --- | --- | --- | --- |
|  |  | No Cefoxitin | 8 mg/L Cefoxitin | No Cefoxitin | 8 mg/L Cefoxitin |
| ***pAmpC* positive and ESBL negative** | 40 | 40 (100%) | 39 (97.5%) | 40 (100%) | 39 (97.5%) |
| *bla*<sub>CMY-2</sub> | 31 | 31 (100%) | 31 (100%) | 31 (100%) | 31 (100%) |
| *bla*<sub>CMY-42</sub> | 3 | 3 (100%) | 3 (100%) | 3 (100%) | 3 (100%) |
| *bla*<sub>CMY-79</sub> | 1 | 1 (100%) | 1 (100%) | 1 (100%) | 1 (100%) |
| *bla*<sub>DHA</sub> | 2 | 2 (100%) | 2 (100%) | 2 (100%) | 2 (100%) |
| *bla*<sub>ACC</sub> | 2 | 2 (100%) | 1 (50%) | 2 (100%) | 1 (50%) |
| *bla*<sub>MOX</sub> | 1 | 1 (100%) | 1 (100%) | 1 (100%) | 1 (100%) |
| ***cAmpC* positive and ESBL negative** | 40 | 40 (100%) | 39 (97.5%) | 40 (100%) | 40 (100%) |
| **ESBL positive and AmpC negative** | 40 | 40 (100%) | 17 (42.5%) | 40 (100%) | 15 (37.5%) |
| *bla*<sub>CTX-M-1</sub> group | 28 | 28 (100%) | 14 (50%) | 28 (100%) | 14 (50%) |
| *bla*<sub>CTX-M-9</sub> group | 12 | 12 (100%) | 3 (25%) | 12 (100%) | 1 (8.3%) |
| **AmpC negative and ESBL negative** | 39 | 1 (2.6%) | 1(2.6%) | 0 (0%) | 0 (0%) |

The cefotaxime + cefoxitin agar showed growth of 39 (97.5%) pAmpC isolates, 39 (97.5%) cAmpC isolates, 17 (42.5%) ESBL isolates and one (2.6%) AmpC/ESBL negative isolate. The ceftazidime + cefoxitin agar showed growth of 39 (97.5%) pAmpC isolates, 40 (100%) cAmpC isolates, 15 (37.5%) ESBL isolates and none of the AmpC/ESBL negative isolates. In both agars the pAmpC isolate that did not grow was the same and appeared to contain a *bla*ACC gene. The ATCC control *E. coli* strain did not grow on any of the evaluated agars.

Table 2 and 3 present the sensitivity and specificity values. Although the agars containing cefoxitin showed a slightly lower sensitivity then the agars without cefoxitin, all confidence intervals overlapped. Specificity was assessed for the 79 AmpC-negative isolates. The agars containing cefoxitin showed a better specificity than the agars without cefoxitin. The specificity of the cefotaxime 1mg/L + cefoxitin 8mg/L and ceftazidime 1mg/L + cefoxitin 8mg/L agar plates were similar, based upon overlap of confidence intervals.

**TABLE 2.** Sensitivity (with 95% confidence intervals) of different agars

|  | pAmpC |  | cAmpC |  |
| --- | --- | --- | --- | --- |
| *Cefotaxime* | 40/40 | 1.00<br>(0.92-1.00) | 40/40 | 1.00<br>(0.92-1.00) |
| *Cefotaxime +*<br>*Cefoxitin* | 39/40 | 0.98<br>(0.86-1.00) | 39/40 | 0.98<br>(0.86-1.00) |
| *Ceftazidime* | 40/40 | 1.00<br>(0.92-1.00) | 40/40 | 1.00<br>(0.92-1.00) |
| *Ceftazidime +*<br>*Cefoxitin* | 39/40 | 0.98<br>(0.86-1.00) | 40/40 | 1.00<br>(0.92-1.00) |

\*Sensitivity based on growth on agar (true positive) divided through all isolates of group (true positive + false negative).

**TABLE 3.**
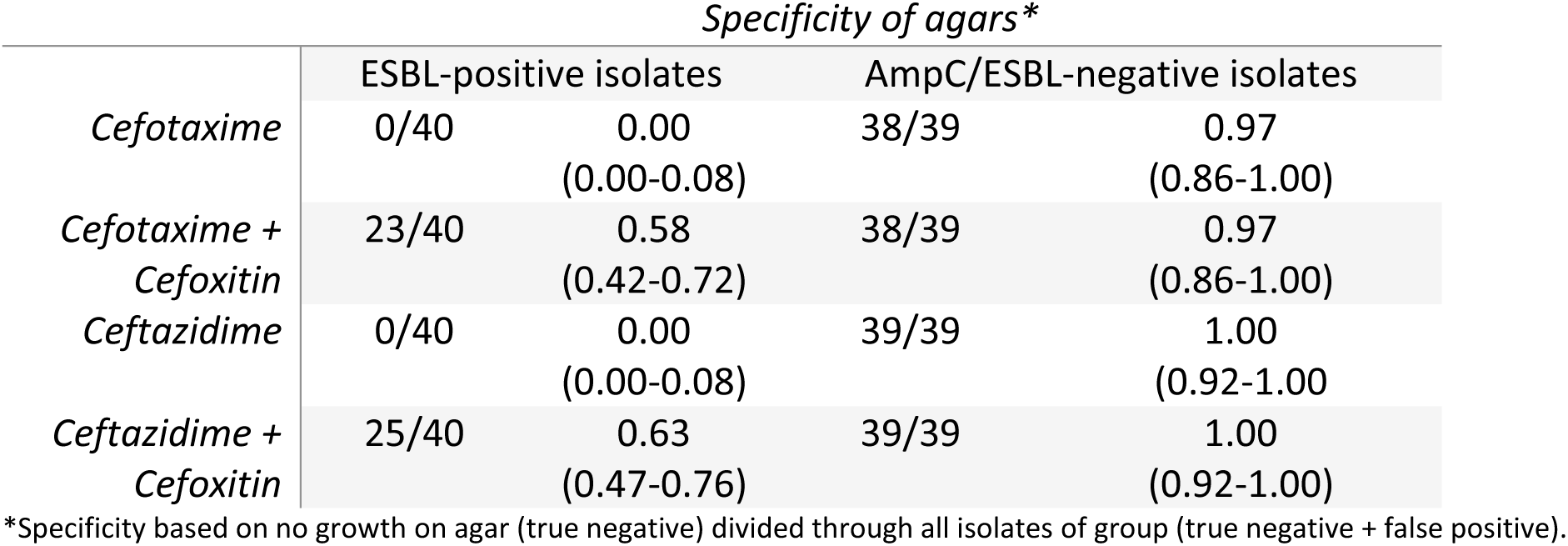
Specificity (with 95% confidence intervals) of different agars

As described above, we found that ESBL isolates had the largest impact on specificity, since ∼40% of them grew on the cefotaxime+cefoxitin or ceftazidime+cefoxitin agars. The prevalence of ESBL in the isolates tested was 25.16%, due to the artificial nature of the isolates chosen. Figs 1 and 2 show the relation between specificity of the cefotaxime 1mg/L + cefoxitin 8mg/L and of the ceftazidime 1mg/L + cefoxitin 8mg/L agar plotted versus ESBL prevalence. An increase in ESBL prevalence in the isolate collection would decrease specificity in a nonlinear function. According to our model a low ESBL prevalence e.g. 5% would lead to a specificity of the cefotaxime 1mg/L + cefoxitin 8mg/L agar of 92% and of the ceftazidime 1mg/L + cefoxitin 8mg/L agar of 95%. A high prevalence of e.g. 50% would decrease specificity of the cefotaxime 1mg/L + cefoxitin 8mg/L agar to 67% and of the ceftazidime 1mg/L + cefoxitin 8mg/L agar to 72%.

**Fig 1.**
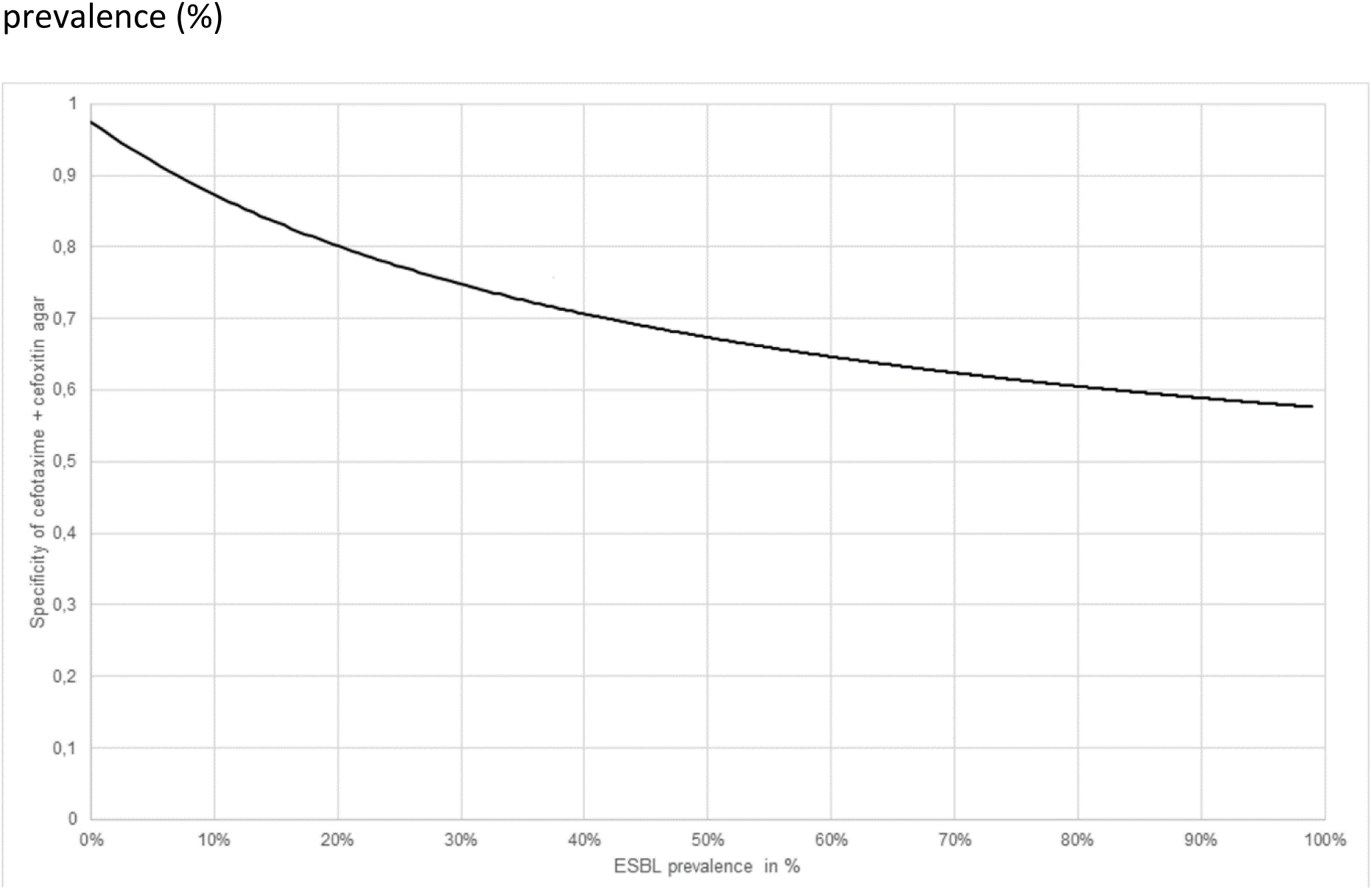
Model of expected specificity of cefotaxime 1mg/L + cefoxitin 8mg/L agar versus ESBL prevalence (%)

**Fig 2.**
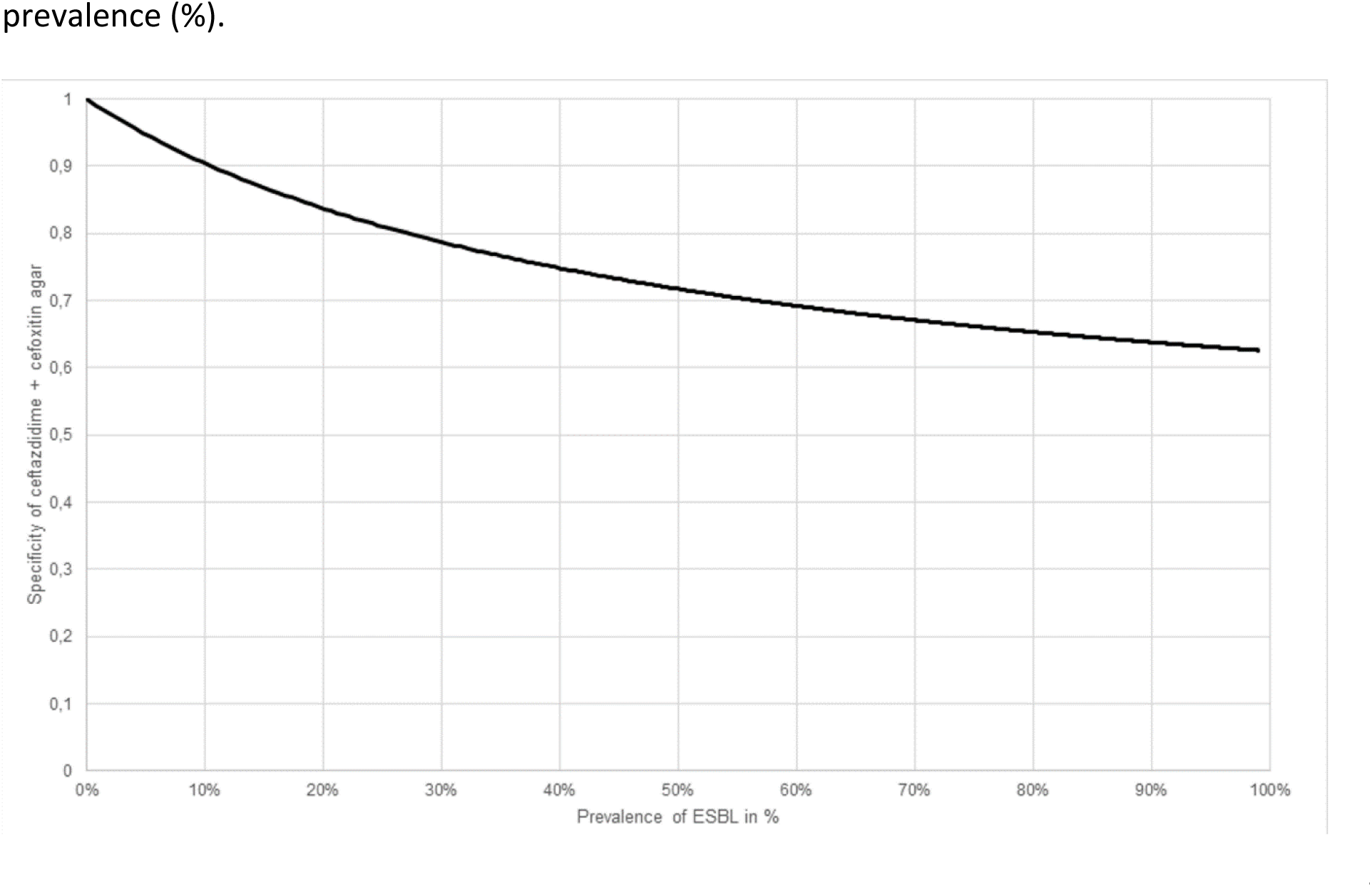
Model of expected specificity of ceftazidime 1mg/L + cefoxitin 8mg/L agar versus ESBL prevalence (%).

## Discussion

Four different screening agars were evaluated on their ability to identify AmpC producing *E. coli*. Cefotaxime- and ceftazidime containing agar showed similar results in sensitivity and specificity for AmpC detection in the tested panel of *E. coli* isolates. The addition of cefoxitin increased specificity for AmpC detection, but did not influence sensitivity.

Although there are no studies on AmpC selective agars, these results are in line with studies showing that cefoxitin might be a useful screening additive for AmpC production(11–13). Polsfuss *et al* analyzed the use of cefoxitin disc diffusion compared to cefotetan for different Enterobacteriales (13). The study included 211 potential AmpC producers based on the basis of cefoxitin inhibition zone diameters of ≤18 mm, cefotetan inhibition zone diameters of ≤16 mm, and/or positive ESBL screening diameters according to CLSI guidelines. The potential AmpC producers were compared to 94 isolates that tested negative in the three mentioned screening criteria. All isolates were subjected to an AmpC multiplex PCR and promoter/attenuator regions of *E. coli* isolates were sequenced. The detection of a pAmpC gene and/or known promoter/attenuator mutations leading to AmpC hyperproduction were considered as gold standard. The majority of acquired AmpC *β*-lactamases were of the CMY-group and DHA-group, none of the strains possessed *β*-lactamases of the ACC, FOX, MOX or ACT/MIR groups. Cefoxitin showed a sensitivity of 97,4% and a specificity of 78,7%. The use of cefotetan, another type of cephamycin, was tested as well, but although this method showed a better specificity (99,3%), sensitivity was much lower (52,6%).

As already described, Reuland *et al* compared different screening methods based upon reduced 3rd generation cephalosporin susceptibility for plasmid-encoded AmpC in Enterobacteriales (14). The study collection consisted out of 356 Enterobacteriales resistant to a third generation cephalosporins (cefotaxime and/or ceftazidime) and/or cefoxitin, of which 68.8% was determined as *E. coli*. A total of 34 pAmpC containing isolates (28 *E. coli*) were detected containing genes from the CMY-group n=29 ; DHA-group n=4 and ACC-group n=1. No analysis on chromosomal AmpC hyperproducers was performed. The strategy using reduced susceptibility to cefotaxime and/or ceftazidime together with reduced susceptibility to cefoxitin showed a sensitivity of 97% and a specificity of 90%. The only isolate not detected contained an ACC-gene.

In our study one isolate containing a ACC-gene did not grow on the cefoxitin containing agars. Although most acquired AmpC *β*-lactamases are inhibited by cephamycines, the ACC-group remains an exception (10). This *β*-lactamase seems to be less common as CMY-group AmpC *β*-lactamases. Nevertheless, ACC-type enzymes have been detected in several countries in Europe (21–23). Outbreaks have been reported as well (24). The use of cefoxitin containing agars may have limitations in areas where the ACC-group is prevalent.

Clearly, there are more limitations in the present study. First, this study is an analytical exploration of sensitivity and specificity based upon retrospectively selected strains. Although the panel of strains was selected on genotype, all strains were obtained from clinical or study collections in which different culture screening criteria were used. We tried to minimize selection bias based upon our screening criteria, but the distribution of our collection may not be completely comparable to a clinical setting. The use of mainly CMY-group pAmpC isolates may have led to overestimation of sensitivity in some degree. However, the CMY-group seems to be the predominant *pampC* gene in clinical settings, and the cefoxitin containing agars may be therefor applicable in these setting for e.g. screening of AmpC rectal carriage.

We remodeled specificity for the cefotaxime 1mg/L + cefoxitin 8mg/L and ceftazidime 1mg/L + cefoxitin 8mg/L agar, as the number of ESBL isolates seems to influence agar specificity. We expect that in a low endemic setting of ESBL specificity of both cefoxitin containing agars will increase. In the Netherlands prevalence of ESBL rectal carriage has been found to range from 5% in a teaching hospital in the South of the Netherlands (25) to 8,6% in general practices in Amsterdam (26). Numbers on rectal carriage in North-America are scarce, but a recent prevalence study on ESBL infections in a hospital in the United States found ranges between 4.7% to 13.4% (27). It is likely that screenings agars in area’s similar to the Dutch setting would have high specificity. In an ESBL high endemic area specificity of the agar will be lower.

Our study was performed using strains cultured in semi-selective pre-enrichment broth. This pre-enrichment step was included to mimic our laboratory algorithm for screening of multiresistant Enterobacterales in rectal samples. Former studies on the screening of third generation cephalosporin resistance have shown that a pre-enrichment strategy results in higher yield compared to direct plating (28–30). We assume that that the use of a pre-enrichment strategy in clinical setting will result in a similar detection range, though inoculum of the original sample may differ. A limit of detection is difficult to obtain, as it is not well known which load in clinical samples is comparable to an *in vitro* setting. Our strategy using the combination of pre-enrichment and selective screening agars was applied in a prevalence study on AmpC in the Amphia hospital (16). Yield of AmpC *E. coli* was comparable to other Dutch prevalence studies (31,32). However, a more elaborate evaluation of clinical samples is needed to confirm our results on sensitivity and specificity of our screening strategy.

In this study the use of screening agars was only assessed on *E. coli,* as this is one of the most prevalent Enterobacteriales with acquired *β*-lactamase genes. We expect the agar to be useful for other AmpC producing Enterobacteriales, however sensitivity and specificity for other species may differ. For example, *K. pneumoniae* without pAmpCs can be cefoxitin-resistant due to loss of porin expression (33). Furthermore, not all existing *β*-lactamase groups were included. No *E. coli* isolates containing AmpC *bla*_FOX_ and *bla*_ACT/MIR_ groups nor the ESBL *bla*_SHV_ and *bla*_TEM_ or other ESBL groups nor carbapenemases were tested. Co-expression of AmpC and ESBL in *E. coli* isolates lacked as well. Although, *bla*_CTX-M_, *bla*_CMY_ and *bla*_DHA_ seem to be most prevalent in the European setting(10,34), future analysis is needed to see if results are applicable to other Enterobacteriales, *β*-lactamase gene groups and *β*-lactamase gene combinations as well.

The AmpC screening agars are not able to differentiate pAmpC vs cAmpC in *E. coli.* Both mechanisms cause elevated 3^rd^ generation cephalosporin MICs, and MIC distributions of cAmpC and pAmpC tend to overlap (35,36).This makes it difficult to create a specific pAmpC or cAmpC screening agar. EUCAST guidelines advise a PCR to distinguish both mechanisms. A screening agar on AmpC may limit the amount strains for molecular confirmation.

## Conclusions

Cefotaxime or ceftazidime had a similar sensitivity and specificity for AmpC in screenings agar in our isolate panel. Addition of cefoxitin to create a more AmpC selective agar had little influence on sensitivity, but increased specificity. Our results apply mainly for *bla*_CMY-type_ producing *E. coli* and further studies are needed to evaluate an agar containing cefotaxime or ceftazidime with cefoxitin in a clinical setting. We expect the screening agars to be feasible to screen for AmpC rectal carriage, when using pre-enrichment similar to current ESBL screening agar strategies. So, agars containing cefotaxime or ceftazidime with cefoxitin do not seem to be inferior in sensitivity compared to agars without cefoxitin, and may be promising when used additional to e.g. a ESBL specific screenings agar for screening of 3^rd^ generation cephalosporin resistance in *E. coli*.

## Supporting information

S 1. Detailed supplementary Materials and Methods

S2. Overview of the panel of E.coli strains

## Abbreviations

ESBL: Extended-spectrum beta-lactamase
ESBL-E: Extended-spectrum beta-lactamase producing Enterobacteriales
pAmpC: plasmid-encoded AmpC
cAmpC: chromosomally-encoded AmpC AmpC-E: AmpC producing Enterobacteriales

## Acknowledgments

The authors thank the laboratory technicians from the laboratory for Microbiology and Infection Control of the Amphia Hospital, the Central Veterinary Institute, the Department of Medical Microbiology, University Medical Center Groningen and the Department of Medical Microbiology, Certe involved in the setup of the strain collection.

## Declarations

### Authors’ contributions

E.D. and J.A.J.W.K. conceived of and designed the study ; J.S. and C.V. were responsible for laboratory processing of the isolates ; J.J.V. conducted molecular analysis of local strains ; K.V., J.W.R., A.M.D.K.S., and M.F.Q.K. provided isolates for the strain collection and revised and approved the manuscript. E.D. performed the analyses and wrote the first draft of the manuscript with input from J.A.J.W.K. All authors contributed to preparation of the final manuscript.

## Additional files

Additional file 1. Detailed supplementary “Materials and Methods”

Additional file 2. Overview of the panel of *E.coli* strains with origin, sequence methode, acquired *β*-lactamases, MLST typing and alterations in promoter and attenuator region

