## Supplementary material for "Detection of AmpC *β*-lactamases in *Escherichia coli* using different screening agars": S 1. Detailed supplementary Materials and Methods

### AmpC *E. coli* isolates from SOM study

In total 27 *E. coli* isolates were selected from the SOM-study (15)based upon the presence of pAmpC encoding genes in the whole genome sequence (WGS) data. In this study perianal swabs taken from hospital patients, were pre-enriched using selective tryptic soy broth (TSB) and subsequently cultured on EbSA screening agar (Cepheid Benelux, Apeldoorn, the Netherlands). Sequencing was performed as described by Kluytmans-van den Bergh *et al* (15). The assembled genomes were uploaded to the online bioinformatics tools ResFinder v2.1 and MLST v1.8 (Center for Genomic Epidemiology, Technical University of Denmark, Lingby, Denmark (version 2.1) (16, 17). ESBL- and pAmpC-encoding genes were reported when at least 60% of the length of the best matching gene in the ResFinder database was covered with a sequence identity of at least 90%. Conventional MLST sequence types were based on the Achtman MLST scheme(18).

In addition, the assembled sequences of each isolate were aligned against the promoter/attenuator region of the cAmpC gene of the *E. coli* K-12 strain MG1655 (GenBank database accession number U00096 (https://www.ncbi.nlm.nih.gov/nuccore/U00096)) using CLC Genomic Workbench version 8.5 (CLC Bio, Qiagen, Hilden,

Germany). The level of hyperproduction of AmpC was based on a former study by Tracz *et al* (7)

**AmpC *E. coli* isolates from Amphia prevalence screening**

In total 22 cAmpC hyperproducing and 4 pAmpC containing *E. coli* isolates were selected from prevalence screening which had been performed in the Amphia hospital described by Den Drijver *et al* (19). Rectal swabs taken from hospital patients were pre-enriched using selective TSB and subsequently cultured on MacConkey agar plate containing cefotaxime (1 mg/L) or MacConkey double agar plate containing cefotaxime (1 mg/L) with cefoxitin (8 mg/L) one side and ceftazidime (1 mg/L) with cefoxitin (8mg/L) other side (Mediaproducts, Groningen, The Netherlands). Presence of pAmpC was analyzed with micro-array MDR CT103 (Check-Points, Wageningen, the Netherlands), alterations in the promoter and attenuator region were analysed with Sanger sequencing as previously described (19).

**pAmpC-encoding clinical *E. coli* isolates from Amphia hospital**

In total 9 pAmpC producing *E. coli* isolates were selected retrospectively from our laboratory database based upon the presence of pAmpC encoding genes. In the laboratory protocol for AmpC screening on clinical samples, *E. coli* isolates with cefotaxime and/or ceftazidime MIC > 1mg/l combined with cefoxitin MIC ≥8mg/l are screened for AmpC production using D68C AmpC & ESBL Detection Set (Mastdiscs, Mastgroup Ltd, Bootle, United Kingdom). The presence of pAmpC genes and/or alterations in the promoter/attenuator region were evaluated similar as in the Amphia prevalence screening described by Den Drijver *et al* (19).

**AmpC *E. coli* isolates from Wageningen University prevalence study**

In total 18 cAmpC hyperproducing *E. coli* isolates were selected retrospectively from a former prevalence study among livestock (e.g. cattle, pigs and broilers) by the Wageningen University.

Screening swabs were cultured on MacConkey agar (product no. 212123, Becton Dickinson) + 1 mg/L cefotaxime (Sigma-Aldrich, Germany)] and inoculated in a selective pre-enrichment broth (Luria–Bertani broth containing 1 mg/L cefotaxime). Phenotypic and genotypic confirmation with micro-array and sanger sequencing were performed similar as described by Dierikx *et al* (20, 21) and Hordijk *et al* (22).

**ESBL-encoding clinical *E. coli* isolates from Amphia hospital**

In total 40 ESBL producing *E. coli* isolates were selected retrospectively from our laboratory database of blood culture *E. coli* isolates containing of CTX-M encoding genes. In the in laboratory protocol for ESBL screening on clinical samples, *E. coli* isolates with cefotaxime and/or ceftazidime MIC > 1mg/l are screened for ESBL production using combination disk diffusion method for cefotaxime, ceftazidime, and cefepime with and without clavulanic acid (Rosco, Taastrup, Denmark) and interpreted according to manufacturer's instructions.

WGS was performed in UMCG using MiSeq (Illumina, San Diego, United States) and assembled with CLC Genomics Workbench 9.0, 9.0.1 or 9.5.2 (Qiagen, Hilden, Germany) as was previously described in more detail by Kluytmans-van den Bergh *et al* (15). Assembly, identification of resistance genes and MLST type and analysis of promoter/attenuator region were performed as described for the SOM study isolates.

**Non-AmpC/non-ESBL producing *E. coli* from STEC-ID-net study**

In total 39 *E. coli* isolates obtained from stools of patients in a former multicentre prospective study, STEC-ID-net, performed from April 2013 to March 2014 in the Dutch regions of Groningen and Rotterdam (23) were selected based on the absence of both AmpC, ESBL genes and known promoter/attenuator mutations related to AmpC hyperproduction. Identification of resistance genes and MLST type and analysis of promoter/attenuator regions were performed as described above (pAmpC SOM study isolates) using the already available WGS data.
