## Supplementary material for "Detection of AmpC *β*-lactamases in *Escherichia coli* using different screening agars": S2. Overview of the panel of E.coli strains

S2. Overview of the panel of *E.coli* strains with origin, sequence methode, acquired β-lactamases, MLST typing and alterations in promoter and attenuator region

| Strain | Group | Study | Host | Sequence method | Acquired β-lactamase | Mulvey type‡ | MLST | -118 | -88 | -82 | -73 | -42 | -32 | -28 | -18 | -16 | -15 | -13 | -11 | -1 | 6 | 11 | 17 | 22 | 24 | 31 | 33 | 34 | 35 | 37 | 58 | 63 | 70 | 81 | cAmpC hyperproduction† |
|---|---|---|---|---|---|---|---|---|---|---|---|---|---|---|---|---|---|---|---|---|---|---|---|---|---|---|---|---|---|---|---|---|---|---|---|
| *K-12 E.coli references strain MG1655 (GenBank database accession number U00096)* | | | | | | | | T | C | A | C | C | T | G | G | – | – | – | C | C | C | | C | C | C | C | G | G | C | G | C | T | C | G | Hyperproduction |
| 1 | pAmpC | SOM-study | human | WGS | CMY-2+TEM wt | 11 | ST4118 | | | | | | | | | | | | | | | | | | | | | | | | | | | | T | | No |
| 2 | pAmpC | SOM-study | human | WGS | CMY-2 | 52 | ST357 | | | | T | | | A | | | | | | | | | | | | | | | | | | | | | | A | No |
| 3 | pAmpC | SOM-study | human | WGS | CMY-2 + TEM wt | 52 | ST131 | | | | T | | | A | | | | | | | | | | | | | | | | | | | | | | A | No |
| 4 | pAmpC | SOM-study | human | WGS | CMY-2 | 52 | ST131 | | | | T | | | A | | | | | | | | | | | | | | | | | | | | | | A | No |
| 5 | pAmpC | SOM-study | human | WGS | CMY-2 | 52 | ST131 | | | | T | | | A | | | | | | | | | | | | | | | | | | | | | | A | No |
| 6 | pAmpC | SOM-study | human | WGS | CMY-2 + TEM wt | 5 | ST648 | | | | T | | | A | | | | | | | | | | | | | | | | | | | T | | | A | No |
| 7 | pAmpC | SOM-study | human | WGS | CMY-2 | 17 | ST457 | | | | T | | | A | | | | | | | | | | | | | | | | | | | | | T | | No |
| 8 | pAmpC | SOM-study | human | WGS | CMY-2 + TEM wt | WT | ST5454 | | | | | | | | | | | | | | | | | | | | | | | | | | | | | | No |
| 9 | pAmpC | SOM-study | human | WGS | CMY-2 | 52 | ST59 | | | | T | | | A | | | | | | | | | | | | | | | | | | | | | | A | No |
| 10 | pAmpC | SOM-study | human | WGS | CMY-2 | 11 | ST38 | | | | | | | | | | | | | | | | | | | | | | | | | | | | T | | No |
| 11 | pAmpC | SOM-study | human | WGS | CMY-2 | 5 | ST1722 | | | | T | | | A | | | | | | | | | | | | | | | | | | | T | | | A | No |
| 12 | pAmpC | SOM-study | human | WGS | CMY-2 + TEM wt + OXA wt | WT | ST1487 | | | | | | | | | | | | | | | | | | | | | | | | | | | | | | No |
| 13 | pAmpC | SOM-study | human | WGS | CMY-2 | 52 | ST131 | | | | T | | | A | | | | | | | | | | | | | | | | | | | | | | A | No |
| 14 | pAmpC | SOM-study | human | WGS | CMY-2 | 52 | ST131 | | | | T | | | A | | | | | | | | | | | | | | | | | | | | | | A | No |
| 15 | pAmpC | SOM-study | human | WGS | CMY-2 | 52 | ST131 | | | | T | | | A | | | | | | | | | | | | | | | | | | | | | | A | No |
| 16 | pAmpC | SOM-study | human | WGS | CMY-2 + TEM wt | 5 | ST1485 | | | | T | | | A | | | | | | | | | | | | | | | | | | | T | | | A | No |
| 17 | pAmpC | SOM-study | human | WGS | CMY-2 + OXA wt | 18 | ST683 | | T | G | | | | | A | | | | | T | | | | | | | | | | | T | | | | No |
| 18 | pAmpC | SOM-study | human | WGS | CMY-2 + TEM wt | 18 | ST1196 | | T | G | | | | | A | | | | | T | | | | | | | | | | | T | | | | No |
| 19 | pAmpC | SOM-study | human | WGS | CMY-2 + OXA wt | 18 | ST23 | | T | G | | | | | A | | | | | T | | | | | | | | | | | T | | | | No |
| 20 | pAmpC | SOM-study | human | WGS | CMY-2 + TEM wt | 18 | ST345 | | T | G | | | | | A | | | | | T | | | | | | | | | | | T | | | | No |
| 21 | pAmpC | SOM-study | human | WGS | CMY-42 | 18 | ST205 | | T | G | | | | | A | | | | | T | | | | | | | | | | | T | | | | No |
| 22 | pAmpC | SOM-study | human | WGS | CMY-42 + TEM wt | WT | ST410 | | | | | | | | | | | | | | | | | | | | | | | | | | | | | | No |
| 23 | pAmpC | SOM-study | human | WGS | CMY-42 | WT | ST410 | | | | | | | | | | | | | | | | | | | | | | | | | | | | | | No |
| 24 | pAmpC | SOM-study | human | WGS | CMY-79 + TEM wt | 18 | ST155 | | T | G | | | | | A | | | | | T | | | | | | | | | | | T | | | | No |
| 25 | pAmpC | SOM-study | human | WGS | DHA + TEM wt | WT | ST1488 | | | | | | | | | | | | | | | | | | | | | | | | | | | | | | No |
| 26 | pAmpC | SOM-study | human | WGS | ACC + TEM wt + OXA wt + SHV wt | 18 | ST224 | | T | G | | | | | A | | | | | T | | | | | | | | | | | T | | | | No |
| 27 | pAmpC | clinical isolate | human | PCR | MOX | WT | unknown | | | | | | | | | | | | | | | | | | | | | | | | | | | | | | No |
| 28 | pAmpC | clinical isolate | human | PCR | CMY-2 | 52 | unknown | | | | T | | | A | | | | | | | | | | | | | | | | | | | | | | A | No |
| 29 | pAmpC | clinical isolate | human | PCR | CMY-2 | 18 | unknown | | T | G | | | | | A | | | | | T | | | | | | | | | | | T | | | | No |
| 30 | pAmpC | clinical isolate | human | PCR | CMY-2 | 18 | unknown | | T | G | | | | | A | | | | | T | | | | | | | | | | | T | | | | No |
| 31 | pAmpC | clinical isolate | human | PCR | CMY-2 | 52 | unknown | | | | T | | | A | | | | | | | | | | | | | | | | | | | | | | A | No |
| 32 | pAmpC | clinical isolate | human | PCR | CMY-2 | WT | unknown | | | | | | | | | | | | | | | | | | | | | | | | | | | | | | No |
| 33 | pAmpC | clinical isolate | human | PCR | CMY-2 | 7 | unknown | | | | T | | | A | | | | | | | | | T | | | | | | | | | | | | | A | No |
| 34 | pAmpC | clinical isolate | human | PCR | CMY-2 | 52 | unknown | | | | T | | | A | | | | | | | | | | | | | | | | | | | | | | A | No |
| 35 | pAmpC | clinical isolate | human | PCR | CMY-2 | unknown | unknown | | | | | | | | | | | | | | | C | | | | | | | | | | | | | | | No |
| 36 | pAmpC | prevalence Amphia | human | PCR | DHA | unknown | unknown | | | | | | | | | | | | | | | | | | | | | | | | T | | | T | | | No |
| 37 | pAmpC | prevalence Amphia | human | PCR | CMY-2 | WT | unknown | | | | | | | | | | | | | | | | | | | | | | | | | | | | | | No |
| 38 | pAmpC | prevalence Amphia | human | PCR | CMY-2 | WT | unknown | | | | | | | | | | | | | | | | | | | | | | | | | | | | | | No |
| 39 | pAmpC | SOM-study | human | WGS | ACC + TEM wt | WT | ST-93 | | | | | | | | | | | | | | | | | | | | | | | | | | | | | | No |
| 40 | pAmpC | prevalence Amphia | human | PCR | CMY-2 | 11 | unknown | | | | | | | | | | | | | | | | | | | | | | | | | | | | T | | No |
| 41 | cAmpC | prevalence Amphia | human | PCR | none | 3 | unknown | | T | G | | T | | | A | | | | | T | | | | | | | | | | | T | | | | Yes (22-fold) |
| 42 | cAmpC | prevalence Amphia | human | PCR | none | 8 | unknown | | | | T | | | | | | | T | | | T | | | | | A | | | | | | | | A | Yes (24-fold) |
| 43 | cAmpC | prevalence Amphia | human | PCR | none | unknown | unknown | | | | | | | | | | | | T | | | | | | | | | del | | | | | T | | Yes ((≈21-68 fold) |
| 44 | cAmpC | prevalence Amphia | human | PCR | none | unknown | unknown | | | | T | | A | | | | | | | | | | | | | | | | | | T | C | | | Yes (8-fold) |
| 45 | cAmpC | prevalence Amphia | human | PCR | none | 3 | unknown | | T | G | | T | | | A | | | | | T | | | | | | | | | | | T | | | | Yes (22-fold) |
| 46 | cAmpC | prevalence Amphia | human | PCR | none | unknown | unknown | | | | | | | | | C | | | | | | | | | | G | | | | | | | T | | Yes ((≈16-64 fold) |
| 47 | cAmpC | prevalence Amphia | human | PCR | none | 3 | unknown | | T | G | | T | | | A | | | | | T | | | | | | | | | | | T | | | | Yes (22-fold) |
| 48 | cAmpC | prevalence Amphia | human | PCR | none | unknown | unknown | | | | T | | A | A | | | | | | | | | | | | | | | | | | | | A | Yes (8-fold) |
| 49 | cAmpC | prevalence WUR | cattle | PCR | none | 3 | unknown | | T | G | | T | | | A | | | | | T | | | | | | | | | | | T | | | | Yes (22-fold) |
| 50 | cAmpC | prevalence WUR | pig | PCR | none | 3 | unknown | | T | G | | T | | | A | | | | | T | | | | | | | | | | | T | | | | Yes (22-fold) |
| 51 | cAmpC | prevalence WUR | broiler | PCR | none | 3 | unknown | | T | G | | T | | | A | | | | | T | | | | | | | | | | | T | | | | Yes (22-fold) |
| 52 | cAmpC | prevalence WUR | cattle | PCR | none | 3 | unknown | | T | G | | T | | | A | | | | | T | | | | | | | | | | | T | | | | Yes (22-fold) |
| 53 | cAmpC | prevalence WUR | cattle | PCR | none | 3 | unknown | | T | G | | T | | | A | | | | | T | | | | | | | | | | | T | | | | Yes (22-fold) |
| 54 | cAmpC | prevalence WUR | pig | PCR | none | 3 | unknown | | T | G | | T | | | A | | | | | T | | | | | | | | | | | T | | | | Yes (22-fold) |
| 55 | cAmpC | prevalence WUR | broiler | PCR | none | 3 | unknown | | T | G | | T | | | A | | | | | T | | | | | | | | | | | T | | | | Yes (22-fold) |
| 56 | cAmpC | prevalence WUR | broiler | PCR | none | 3 | unknown | | T | G | | T | | | A | | | | | T | | | | | | | | | | | T | | | | Yes (22-fold) |
| 57 | cAmpC | prevalence WUR | broiler | PCR | TEM wt | 3 | unknown | | T | G | | T | | | A | | | | | T | | | | | | | | | | | T | | | | Yes (22-fold) |
| 58 | cAmpC | prevalence WUR | broiler | PCR | none | 3 | unknown | | T | G | | T | | | A | | | | | T | | | | | | | | | | | T | | | | Yes (22-fold) |
| 59 | cAmpC | prevalence WUR | cattle | PCR | TEM wt | 3 | unknown | | T | G | | T | | | A | | | | | T | | | | | | | | | | | T | | | | Yes (22-fold) |
| 60 | cAmpC | prevalence WUR | pig | PCR | none | 3 | unknown | | T | G | | T | | | A | | | | | T | | | | | | | | | | | T | | | | Yes (22-fold) |
| 61 | cAmpC | prevalence WUR | pig | PCR | TEM wt | 3 | unknown | | T | G | | T | | | A | | | | | T | | | | | | | | | | | T | | | | Yes (22-fold) |
| 62 | cAmpC | prevalence WUR | cattle | PCR | none | 3 | unknown | | T | G | | T | | | A | | | | | T | | | | | | | | | | | T | | | | Yes (22-fold) |
| 63 | cAmpC | prevalence WUR | pig | PCR | TEM wt | 3 | unknown | | T | G | | T | | | A | | | | | T | | | | | | | | | | | T | | | | Yes (22-fold) |
| 64 | cAmpC | prevalence WUR | pig | PCR | none | 3 | unknown | | T | G | | T | | | A | | | | | T | | | | | | | | | | | T | | | | Yes (22-fold) |
| 65 | cAmpC | prevalence Amphia | human | PCR | none | 23 | unknown* | | | | T | | A | | | | | | | | | | | | | | | | | | | | | A | Yes (19-fold) |
| 66 | cAmpC | prevalence Amphia | human | PCR | none | 8 | unknown* | | | | T | | | | | | | T | | | T | | | | | A | | | | | | | | A | Yes (24-fold) |
| 67 | cAmpC | prevalence Amphia | human | PCR | none | 3 | unknown* | | T | G | | T | | | A | | | | | T | | | | | | | | | | | T | | | | Yes (22-fold) |
| 68 | cAmpC | prevalence Amphia | human | PCR | none | unknown | unknown | | T | G | | T | | | A | | | | | T | | | | A | | | | | | | T | | | | Yes (≈60-140-fold) |
| 69 | cAmpC | prevalence Amphia | human | PCR | TEM wt | 3 | unknown* | | T | G | | T | | | A | | | | | T | | | | | | | | | | | T | | | | Yes (22-fold) |
| 70 | cAmpC | prevalence Amphia | human | PCR | none | 3 | unknown | | T | G | | T | | | A | | | | | T | | | | | | | | | | | T | | | | Yes (22-fold) |
| 71 | cAmpC | prevalence Amphia | human | PCR | none | 2 | unknown* | | | | T | | A | | | | | | | | | | | | | | | | | | T | C | | A | Yes (10-fold) |
| 72 | cAmpC | prevalence Amphia | human | PCR | none | unknown | unknown | | | | T | | | | | | | T | | | | | | | | | A | | | | T | C | | A | Yes (24-fold) |
| 73 | cAmpC | prevalence Amphia | human | PCR | none | 3 | unknown | | T | G | | T | | | A | | | | | T | | | | | | | | | | | T | | | | Yes (22-fold) |
| 74 | cAmpC | prevalence Amphia | human | PCR | none | 3 | unknown | | T | G | | T | | | A | | | | | T | | | | | | | | | | | T | | | | Yes (22-fold) |
| 75 | cAmpC | prevalence Amphia | human | PCR | none | 3 | unknown | | T | G | | T | | | A | | | | | T | | | | | | | | | | | T | | | | Yes (22-fold) |
| 76 | cAmpC | prevalence Amphia | human | PCR | none | 23 | unknown | | | | T | | A | | | | | | | | | | | | | | | | A | | | | | | Yes (19-fold) |
| 77 | cAmpC | prevalence Amphia | human | PCR | none | 3 | unknown | | T | G | | T | | | A | | | | | T | | | | | | | | | | | T | | | | Yes (22-fold) |
| 78 | cAmpC | prevalence WUR | cattle | PCR | none | 3 | unknown | | T | G | | T | | | A | | | | | T | | | | | | | | | | | T | | | | Yes (22-fold) |
| 79 | cAmpC | prevalence WUR | pig | PCR | none | 3 | unknown | | T | G | | T | | | A | | | | | T | | | | | | | | | | | T | | | | Yes (22-fold) |
| 80 | cAmpC | prevalence Amphia | human | PCR | none | 3 | unknown | | T | G | | T | | | A | | | | | T | | | | | | | | | | | T | | | | Yes (22-fold) |
| 81 | ESBL | clinical isolate | human | WGS | CTX-M-9 | 11 | ST38 | | | | | | | | | | | | | | | | | | | | | | | | | | | | T | | No |
| 82 | ESBL | clinical isolate | human | WGS | CTX-M-14 | 52 | ST7092 | | | | T | | | A | | | | | | | | | | | | | | | | | | | | | | A | No |
| 83 | ESBL | clinical isolate | human | WGS | CTX-M-27 | 11 | ST38 | | | | | | | | | | | | | | | | | | | | | | | | | | | | T | | No |
| 84 | ESBL | clinical isolate | human | WGS | CTX-M-15 | 52 | ST131 | | | | T | | | A | | | | | | | | | | | | | | | | | | | | | | A | No |
| 85 | ESBL | clinical isolate | human | WGS | CTX-M-15 | WT | ST410 | | | | | | | | | | | | | | | | | | | | | | | | | | | | | | No |
| 86 | ESBL | clinical isolate | human | WGS | CTX-M-15 | 52 | ST131 | | | | T | | | A | | | | | | | | | | | | | | | | | | | | | | A | No |
| 87 | ESBL | clinical isolate | human | WGS | CTX-M-15 | 52 | ST131 | | | | T | | | A | | | | | | | | | | | | | | | | | | | | | | A | No |
| 88 | ESBL | clinical isolate | human | WGS | CTX-M-15 | 52 | ST131 | | | | T | | | A | | | | | | | | | | | | | | | | | | | | | | A | No |
| 89 | ESBL | clinical isolate | human | WGS | CTX-M-1 | 52 | ST131 | | | | T | | | A | | | | | | | | | | | | | | | | | | | | | | A | No |
| 90 | ESBL | clinical isolate | human | WGS | CTX-M-15 | 11 | ST38 | | | | | | | | | | | | | | | | | | | | | | | | | | | | T | | No |
| 91 | ESBL | clinical isolate | human | WGS | CTX-M-15 | 52 | ST131-like | | | | T | | | A | | | | | | | | | | | | | | | | | | | | | | A | No |
| 92 | ESBL | clinical isolate | human | WGS | CTX-M-15 | 52 | ST131 | | | | T | | | A | | | | | | | | | | | | | | | | | | | | | | A | No |
| 93 | ESBL | clinical isolate | human | WGS | CTX-M-15 | 52 | ST131 | | | | T | | | A | | | | | | | | | | | | | | | | | | | | | | A | No |
| 94 | ESBL | clinical isolate | human | WGS | CTX-M-15 | 52 | ST131 | | | | T | | | A | | | | | | | | | | | | | | | | | | | | | | A | No |
| 95 | ESBL | clinical isolate | human | WGS | CTX-M-55 | 52 | ST131 | | | | T | | | A | | | | | | | | | | | | | | | | | | | | | | A | No |
| 96 | ESBL | clinical isolate | human | WGS | CTX-M-15 | 52 | ST131 | | | | T | | | A | | | | | | | | | | | | | | | | | | | | | | A | No |
| 97 | ESBL | clinical isolate | human | WGS | CTX-M-15 | 11 | ST315 | | | | | | | | | | | | | | | | | | | | | | | | | | | | T | | No |
| 98 | ESBL | clinical isolate | human | WGS | CTX-M-15 | WT | ST617 | | | | | | | | | | | | | | | | | | | | | | | | | | | | | | No |
| 99 | ESBL | clinical isolate | human | WGS | CTX-M-15 | 45 | ST73 | | | | T | | | | | | | | | | | | | | | | | | | | | | | | | A | No |
| 100 | ESBL | clinical isolate | human | WGS | CTX-M-27 | 52 | ST131 | | | | T | | | A | | | | | | | | | | | | | | | | | | | | | | A | No |
| 101 | ESBL | clinical isolate | human | WGS | CTX-M-1 | 52 | ST357 | | | | T | | | A | | | | | | | | | | | | | | | | | | | | | | A | No |
| 102 | ESBL | clinical isolate | human | WGS | CTX-M-15 | 52 | ST131 | | | | T | | | A | | | | | | | | | | | | | | | | | | | | | | A | No |
| 103 | ESBL | clinical isolate | human | WGS | CTX-M-1 | WT | ST10 | | | | | | | | | | | | | | | | | | | | | | | | | | | | | | No |
| 104 | ESBL | clinical isolate | human | WGS | CTX-M-15 | 52 | ST131 | | | | T | | | A | | | | | | | | | | | | | | | | | | | | | | A | No |
| 105 | ESBL | clinical isolate | human | WGS | CTX-M-1 | 5 | ST648 | | | | T | | | A | | | | | | | | | | | | | | | | | | | T | | | A | No |
| 106 | ESBL | clinical isolate | human | WGS | CTX-M-15 | 52 | ST131 | | | | T | | | A | | | | | | | | | | | | | | | | | | | | | | A | No |
| 107 | ESBL | clinical isolate | human | WGS | CTX-M-15 | 11 | ST501 | | | | | | | | | | | | | | | | | | | | | | | | | | | | T | | No |
| 108 | ESBL | clinical isolate | human | WGS | CTX-M-15 | 52 | ST131 | | | | T | | | A | | | | | | | | | | | | | | | | | | | | | | A | No |
| 109 | ESBL | clinical isolate | human | WGS | CTX-M-14 | 52 | ST131 | | | | T | | | A | | | | | | | | | | | | | | | | | | | | | | A | No |
| 110 | ESBL | clinical isolate | human | WGS | CTX-M-14 | 12 | ST12 | | | | T | | | | | | | | | | | | | | | | | | | | T | C | | A | No |
| 111 | ESBL | clinical isolate | human | WGS | CTX-M-15 | WT | ST410 | | | | | | | | | | | | | | | | | | | | | | | | | | | | | | No |
| 112 | ESBL | clinical isolate | human | WGS | CTX-M-1 | WT | ST10 | | | | | | | | | | | | | | | | | | | | | | | | | | | | | | No |
| 113 | ESBL | clinical isolate | human | WGS | CTX-M-14 | 11 | ST69 | | | | | | | | | | | | | | | | | | | | | | | | | | | | T | | No |
| 114 | ESBL | clinical isolate | human | WGS | CTX-M-14 | 5 | ST354 | | | | T | | | A | | | | | | | | | | | | | | | | | | | T | | | A | No |
| 115 | ESBL | clinical isolate | human | WGS | CTX-M-15 | 52 | ST131 | | | | T | | | A | | | | | | | | | | | | | | | | | | | | | | A | No |
| 116 | ESBL | clinical isolate | human | WGS | CTX-M-14b | unknown | ST349 | | | | | | | | | | | | | | | | | | | | | | | | T | | | T | | | No |
| 117 | ESBL | clinical isolate | human | WGS | CTX-M-27 | 52 | ST131 | | | | T | | | A | | | | | | | | | | | | | | | | | | | | | | A | No |
| 118 | ESBL | clinical isolate | human | WGS | CTX-M-15 | 52 | ST131 | | | | T | | | A | | | | | | | | | | | | | | | | | | | | | | A | No |
| 119 | ESBL | clinical isolate | human | WGS | CTX-M-14b | 11 | ST38 | | | | | | | | | | | | | | | | | | | | | | | | | | | | T | | No |
| 120 | ESBL | clinical isolate | human | WGS | CTX-M-27 | 52 | ST131 | | | | T | | | A | | | | | | | | | | | | | | | | | | | | | | A | No |
| 121 | non-AmpC/non-ESBL | STEC/EIEC UMCG | human | WGS | none | 18 | ST-677 | | T | G | | | | | A | | | | | T | | | | | | | | | | | T | | | | No |
| 122 | non-AmpC/non-ESBL | STEC/EIEC UMCG | human | WGS | none | WT | ST-548 | | | | | | | | | | | | | | | | | | | | | | | | | | | | | | No |
| 123 | non-AmpC/non-ESBL | STEC/EIEC UMCG | human | WGS | none | 18 | ST-300 | | T | G | | | | | A | | | | | T | | | | | | | | | | | T | | | | No |
| 124 | non-AmpC/non-ESBL | STEC/EIEC UMCG | human | WGS | none | 18 | ST-21 | | T | G | | | | | A | | | | | T | | | | | | | | | | | T | | | | No |
| 125 | non-AmpC/non-ESBL | STEC/EIEC UMCG | human | WGS | none | 18 | ST-13 | | T | G | | | | | A | | | | | T | | | | | | | | | | | T | | | | No |
| 126 | non-AmpC/non-ESBL | STEC/EIEC UMCG | human | WGS | none | 18 | ST-33 | | T | G | | | | | A | | | | | T | | | | | | | | | | | T | | | | No |
| 127 | non-AmpC/non-ESBL | STEC/EIEC UMCG | human | WGS | none | 18 | ST-17 | | T | G | | | | | A | | | | | T | | | | | | | | | | | T | | | | No |
| 128 | non-AmpC/non-ESBL | STEC/EIEC UMCG | human | WGS | none | 12 | ST-583 | | | | T | | | | | | | | | | | | | | | | | | | | T | C | | A | No |
| 129 | non-AmpC/non-ESBL | STEC/EIEC UMCG | human | WGS | none | 18 | ST-21 | | T | G | | | | | A | | | | | T | | | | | | | | | | | T | | | | No |
| 130 | non-AmpC/non-ESBL | STEC/EIEC UMCG | human | WGS | none | 18 | ST-442 | | T | G | | | | | A | | | | | T | | | | | | | | | | | T | | | | No |
| 131 | non-AmpC/non-ESBL | STEC/EIEC UMCG | human | WGS | none | 18 | ST-33 | | T | G | | | | | A | | | | | T | | | | | | | | | | | T | | | | No |
| 132 | non-AmpC/non-ESBL | STEC/EIEC UMCG | human | WGS | none | WT | ST-753 | | | | | | | | | | | | | | | | | | | | | | | | | | | | | | No |
| 133 | non-AmpC/non-ESBL | STEC/EIEC UMCG | human | WGS | none | 18 | ST-447 | | T | G | | | | | A | | | | | T | | | | | | | | | | | T | | | | No |
| 134 | non-AmpC/non-ESBL | STEC/EIEC UMCG | human | WGS | none | WT | ST-32 | | | | | | | | | | | | | | | | | | | | | | | | | | | | | | No |
| 135 | non-AmpC/non-ESBL | STEC/EIEC UMCG | human | WGS | none | 18 | ST-25 | | T | G | | | | | A | | | | | T | | | | | | | | | | | T | | | | No |
| 136 | non-AmpC/non-ESBL | STEC/EIEC UMCG | human | WGS | none | 18 | ST-21 | | T | G | | | | | A | | | | | T | | | | | | | | | | | T | | | | No |
| 137 | non-AmpC/non-ESBL | STEC/EIEC UMCG | human | WGS | none | 18 | ST-21 | | T | G | | | | | A | | | | | T | | | | | | | | | | | T | | | | No |
| 138 | non-AmpC/non-ESBL | STEC/EIEC UMCG | human | WGS | none | 12 | ST-6045 | | | | T | | | | | | | | | | | | | | | | | | | | T | C | | A | No |
| 139 | non-AmpC/non-ESBL | STEC/EIEC UMCG | human | WGS | none | 18 | ST-29 | | T | G | | | | | A | | | | | T | | | | | | | | | | | T | | | | No |
| 140 | non-AmpC/non-ESBL | STEC/EIEC UMCG | human | WGS | none | 18 | ST-21 | | T | G | | | | | A | | | | | T | | | | | | | | | | | T | | | | No |
| 141 | non-AmpC/non-ESBL | STEC/EIEC UMCG | human | WGS | none | 18 | ST-655 | | T | G | | | | | A | | | | | T | | | | | | | | | | | T | | | | No |
| 142 | non-AmpC/non-ESBL | STEC/EIEC UMCG | human | WGS | none | WT | ST-10 | | | | | | | | | | | | | | | | | | | | | | | | | | | | | | No |
| 143 | non-AmpC/non-ESBL | STEC/EIEC UMCG | human | WGS | none | 18 | ST-20 | | T | G | | | | | A | | | | | T | | | | | | | | | | | T | | | | No |
| 144 | non-AmpC/non-ESBL | STEC/EIEC UMCG | human | WGS | none | WT | ST-11 | | | | | | | | | | | | | | | | | | | | | | | | | | | | | | No |
| 145 | non-AmpC/non-ESBL | STEC/EIEC UMCG | human | WGS | none | WT | ST-11 | | | | | | | | | | | | | | | | | | | | | | | | | | | | | | No |
| 146 | non-AmpC/non-ESBL | STEC/EIEC UMCG | human | WGS | none | WT | ST-10 | | | | | | | | | | | | | | | | | | | | | | | | | | | | | | No |
| 147 | non-AmpC/non-ESBL | STEC/EIEC UMCG | human | WGS | none | WT | ST-10 | | | | | | | | | | | | | | | | | | | | | | | | | | | | | | No |
| 148 | non-AmpC/non-ESBL | STEC/EIEC UMCG | human | WGS | none | WT | ST-10 | | | | | | | | | | | | | | | | | | | | | | | | | | | | | | No |
| 149 | non-AmpC/non-ESBL | STEC/EIEC UMCG | human | WGS | TEM wt | 12 | ST-4554 | | | | T | | | | | | | | | | | | | | | | | | | | T | C | | A | No |
| 150 | non-AmpC/non-ESBL | STEC/EIEC UMCG | human | WGS | none | 18 | ST-2178 | | T | G | | | | | A | | | | | T | | | | | | | | | | | T | | | | No |
| 151 | non-AmpC/non-ESBL | STEC/EIEC UMCG | human | WGS | none | WT | ST-10 | | | | | | | | | | | | | | | | | | | | | | | | | | | | | | No |
| 152 | non-AmpC/non-ESBL | STEC/EIEC UMCG | human | WGS | none | WT | ST-10 | | | | | | | | | | | | | | | | | | | | | | | | | | | | | | No |
| 153 | non-AmpC/non-ESBL | STEC/EIEC UMCG | human | WGS | none | WT | ST-10 | | | | | | | | | | | | | | | | | | | | | | | | | | | | | | No |
| 154 | non-AmpC/non-ESBL | STEC/EIEC UMCG | human | WGS | none | WT | ST-10 | | | | | | | | | | | | | | | | | | | | | | | | | | | | | | No |
| 155 | non-AmpC/non-ESBL | STEC/EIEC UMCG | human | WGS | none | WT | ST-11 | | | | | | | | | | | | | | | | | | | | | | | | | | | | | | No |
| 156 | non-AmpC/non-ESBL | STEC/EIEC UMCG | human | WGS | none | WT | ST-11 | | | | | | | | | | | | | | | | | | | | | | | | | | | | | | No |
| 157 | non-AmpC/non-ESBL | STEC/EIEC UMCG | human | WGS | none | 18 | ST-17 | | T | G | | | | | A | | | | | T | | | | | | | | | | | T | | | | No |
| 158 | non-AmpC/non-ESBL | STEC/EIEC UMCG | human | WGS | none | 18 | ST-17 | | T | G | | | | | A | | | | | T | | | | | | | | | | | T | | | | No |
| 159 | non-AmpC/non-ESBL | STEC/EIEC UMCG | human | WGS | none | 18 | ST-33 | | T | G | | | | | A | | | | | T | | | | | | | | | | | T | | | | No |
| 160 | non-AmpC/non-ESBL | ATCC Castellani and Chalmers 25922 | human | PCR | none | 45 | ST-73 | | | | T | | | | | | | | | | | | | | | | | | | | | | | | | A | No |

-35 box    -10 box    attenuator

displaced -35 box    displaced -10 box

‡ As defined by Mulvey et al. (2005). The reader should refer to Table 3 of the given reference to see a complete characterization of each promoter type.

† Based upon expected hyperproduction defined by Tracz et al. (2007 [16])
